# Integrating crop growth models with whole genome prediction through approximate Bayesian computation

**DOI:** 10.1101/014100

**Authors:** Frank Technow, Carlos D. Messina, L. Radu Totir, Mark Cooper

## Abstract

Genomic selection, enabled by whole genome prediction (WGP) methods, is revolutionizing plant breeding. Existing WGP methods have been shown to deliver accurate predictions in the most common settings, such as prediction of across environment performance for traits with additive gene effects. However, prediction of traits with non-additive gene effects and prediction of genotype by environment interaction (G×E), continues to be challenging. Previous attempts to increase prediction accuracy for these particularly difficult tasks employed prediction methods that are purely statistical in nature. Augmenting the statistical methods with biological knowledge has been largely overlooked thus far. Crop growth models (CGMs) attempt to represent the impact of functional relationships between plant physiology and the environment in the formation of yield and similar output traits of interest. Thus, they can explain the impact of G×E and certain types of non-additive gene effects on the expressed phenotype. *Approximate Bayesian computation* (ABC), a novel and powerful computational procedure, allows the incorporation of CGMs directly into the estimation of whole genome marker effects in WGP. Here we provide a proof of concept study for this novel approach and demonstrate its use with synthetic data sets. We show that this novel approach can be considerably more accurate than the benchmark WGP method GBLUP in predicting performance in environments represented in the estimation set as well as in previously unobserved environments for traits determined by non-additive gene effects. We conclude that this proof of concept demonstrates that using ABC for incorporating biological knowledge in the form of CGMs into WGP is a very promising and novel approach to improving prediction accuracy for some of the most challenging scenarios in plant breeding and applied genetics.

## Introduction

Genomic selection (Meuwissen et al. [1]), enabled by whole genome prediction (WGP) methods, is revolutionizing plant breeding (Cooper et al. [2]). Since its inception, attempts to improve prediction accuracy have focused on: developing improved and specialized statistical models (Yang and Tempelman [3], Heslot et al. [4], Kärkkäinen and Sillanpää [5], Technow and Melchinger [6]), increasing the marker density used (Meuwissen and Goddard [7], Erbe et al. [8], Ober et al. [9]), increasing the size and defining optimal designs of estimation sets (Rincent et al. [10], Windhausen et al. [11], Technow et al. [12], Hickey et al. [13]) and better understanding the genetic determinants driving prediction accuracy (Daetwyler et al. [14], Habier et al. [15]).

In-silico phenotypic prediction, enabled by dynamic crop growth models (CGMs), dates back to the late 1960’s (van Ittersum et al. [16]) and it has constantly evolved through inclusion of scientific advances made in plant physiology, soil science and micrometeorology (Keating et al. [17]; van Ittersum et al. [16]). CGMs used in plant breeding are structured around concepts of resource capture, utilization efficiency and allocation among plant organs (Cooper et al. [18]; Hammer et al. [19]; Passioura [20]; Yin et al. [21]) and are used to: characterize environments (Chapman et al. [22]; Löffler et al [23]), predict consequences of trait variation on yield within a genotype x environment x management context (Hammer et al. [24]), evaluate breeding strategies (Chapman et al. [25]; Messina et al. [26]; Messina et al. [27]), and assess hybrid performance (Cooper et al. [2]).

Early attempts to extend the use of CGMs to enable genetic prediction have focused on developing genetic models for parameters of main process equations within the CGM (Chenu et al. [28]; Messina et al. [29]; Yin et al. [21]). Linking quantitative trait locus (QTL) models and CGMs for complex traits motivated adapting CGMs to improve the connectivity between physiology and genetics of the adaptive traits (Hammer et al. [30]; Messina et al. [27]; Yin et al. [21]). However, despite a tremendous body of knowledge and experience, CGMs were largely ignored for the purpose of WGP.

There is ample evidence for the importance of epistasis in crops, including for economically important traits such as grain yield in maize (Wolf and Hallauer [31], Eta-Ndu and Openshaw [32]; Holland [33]). Yield and other complex traits are the product of intricate interactions between component traits on lower hierarchical levels (Cooper et al. [34]; Hammer et al. [19]; Riedelsheimer et al. [35]). If the relationship among the underlying component traits is nonlinear, epistatic effects can occur on the phenotypic level of complex traits even if the gene action is purely additive when characterized at the level of the component traits (Holland [33]). This phenomenon was first described for multiplicative relationships among traits by Richey [36] and later quantified by Melchinger et al. [37]. CGMs, which explicitly model these nonlinear relationships among traits, have therefore the potential to open up novel avenues towards accounting for epistatic effects in WGP models by explicit incorporation of biological knowledge.

The target population of environments for plant breeding programs is subject to continuous re-evaluation (Cooper et al. [2]). To select for performance in specific environments, genotype by environment (G×E) interactions have to be predicted. Genomic prediction of G×E interactions is therefore of great interest for practical applications of breeding theory. Previous attempts incorporated G×E interactions in WGP models through environment specific marker effects (Schulz-Streeck et al. [38]) or genetic and environmental covariances (Burgeño et al. [39]). Later Jarquín et al. [40] and Heslot et al. [41] developed WGP models that accounted for G×E interactions by means of environmental covariates.

While these previous attempts are promising, they are purely statistical in nature and do not leverage the substantial biological insights into the mechanisms determining performance in specific environments. CGMs are an embodiment of this biological knowledge and might serve as a key component in novel WGP models for predicting G×E interactions. In fact, Heslot et al. [41] recognized this potential for CGMs. However, they employed them only for computing stress covariates from environmental data, which were subsequently used as covariates in purely statistical WGP models.

Given the potential merits of integrating CGMs in WGP, the question arises of how to combine the two in a unified predictive system. The ever increasing computational power of modern computing environments allows for efficient simulation from the most complex of models, such as CGMs (Messina et al. [27]). This computational power is leveraged by *approximate Bayesian computation* (ABC) methods, which replace the calculation of a likelihood function with a simulation step, and thereby facilitate analysis when calculation of a likelihood function is impossible or computationally prohibitive. ABC methods were developed in population genetics, where they helped solve otherwise intractable problems (Tavare et al. [42]; Pritchard et al. [43]; Csilléry et al. [44]; Lopes and Beaumont [45]). However, ABC methods were rapidly adopted in other scientific fields, such as ecology (Lawson Handley et al. [46]), systems biology (Liepe et al. [47]) and hydrology (Sadegh and Vrugt [48]). Recently, Marjoram et al. [49] proposed using ABC methods for incorporating the biological knowledge represented in gene regulatory networks into genome-wide association studies, arguing that this might present a solution to the “missing heritability” problem.

Here we make the case that ABC may hold great promise for enabling novel approaches to WGP as well. Thus, the objective of this study is to provide a proof of concept, based on synthetic data sets, for using ABC as a mechanism for incorporating the substantial biological knowledge embodied in CGMs into a novel WGP approach.

## Materials and Methods

### CGM and environmental data

We used the maize CGM developed by Muchow et al. [50], which models maize grain yield development as a function of plant population (plants *m*^−2^), daily temperature (°*C*) and solar radiation (*M J m*^−2^) as well as several genotype dependent physiological traits. These traits were total leaf number (TLN), area of largest leaf (AM), solar radiation use efficiency (SRE) and thermal units to physiological maturity (MTU). Details on the calculation of trait values for the genotypes in the synthetic data set are provided later. However, the values used were within typical ranges reported in the literature. The simulated intervals for TLN, AM, SRE and MTU were [6, 23] (Meghji et al. [51], Muchow et al. [50]), [700, 800] (Muchow et al. [50], Elings [52]), [1.5, 1.7] (Muchow and Davis [53] and [1050, 1250] (McGarrahan and Dale [54], Muchow [55], Nielsen et al. [56]), respectively, with average values at the midpoints of the intervals.

We chose Champaign/Illinois (40.08° N, 88.24° W) as a representative US Corn Belt location. Temperature and solar radiation data were obtained for the years 2012 and 2013 (Data provided by the Water and Atmospheric Resources Monitoring Program, a part of the Illinois State Water Survey (ISWS) located in Champaign and Peoria, Illinois, and on the web at www.sws.uiuc.edu/warm). The sowing date in 2012 was April 15th and in 2013 it was May 15th. We modified the original CGM of Muchow et al. [50] by enforcing a maxium length of the growing season, after which crop growth simulation was terminated, regardless of wether the genotype reached full physiological maturity or not. The length of the growing season in 2012 was 120 days from sowing and in 2013 it was 130 days from sowing. Both durations are within the range typically observed in the US Corn Belt (Neild and Newman [57]). In 2012 the plant population was 8 plants *m*^−2^ and in 2013 the plant population was 10 plants *m*^−2^. The 2012 and 2013 environments therefore differed not only in temperature and solar radiation but also in management practices. The temperature and solar radiation from date of sowing is shown in Fig. 1. Typical total biomass and grain yield development curves for early, intermediate and late maturing genotypes in the 2012 and 2013 environments are shown in Fig. 2 and corresponding curves for development of total and senescent leaf area in Fig. S1.

**Figure 1.**
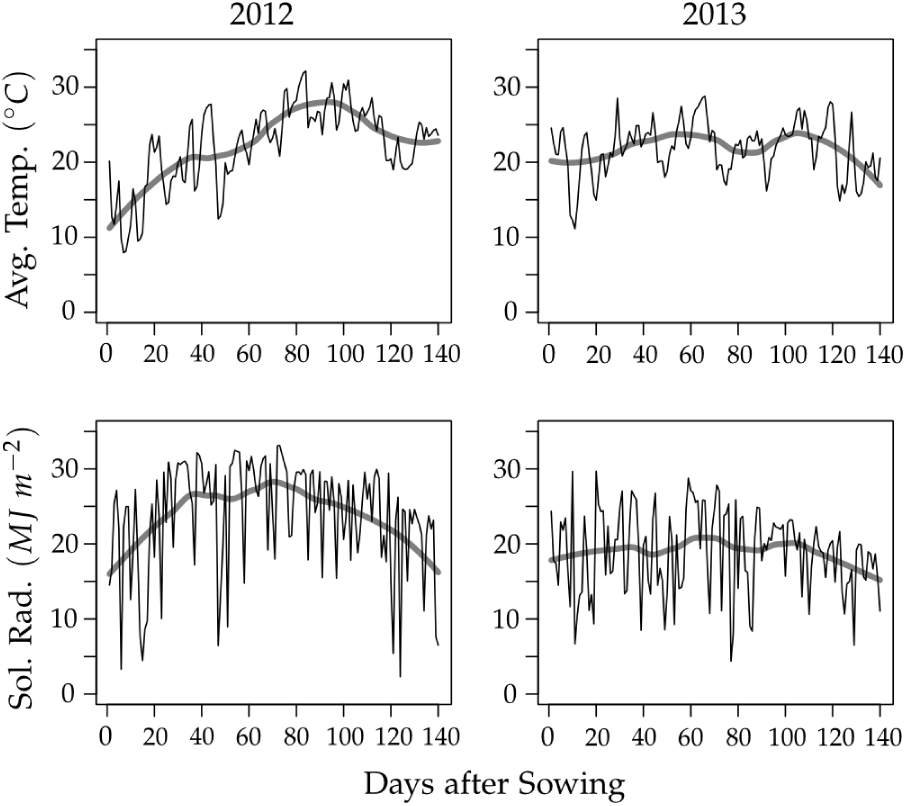
Daily average temperature and solar radiation at Champaign, Illinois in 2012 and 2013. The thick grey line shows a smoothed curve.

**Figure 2.**
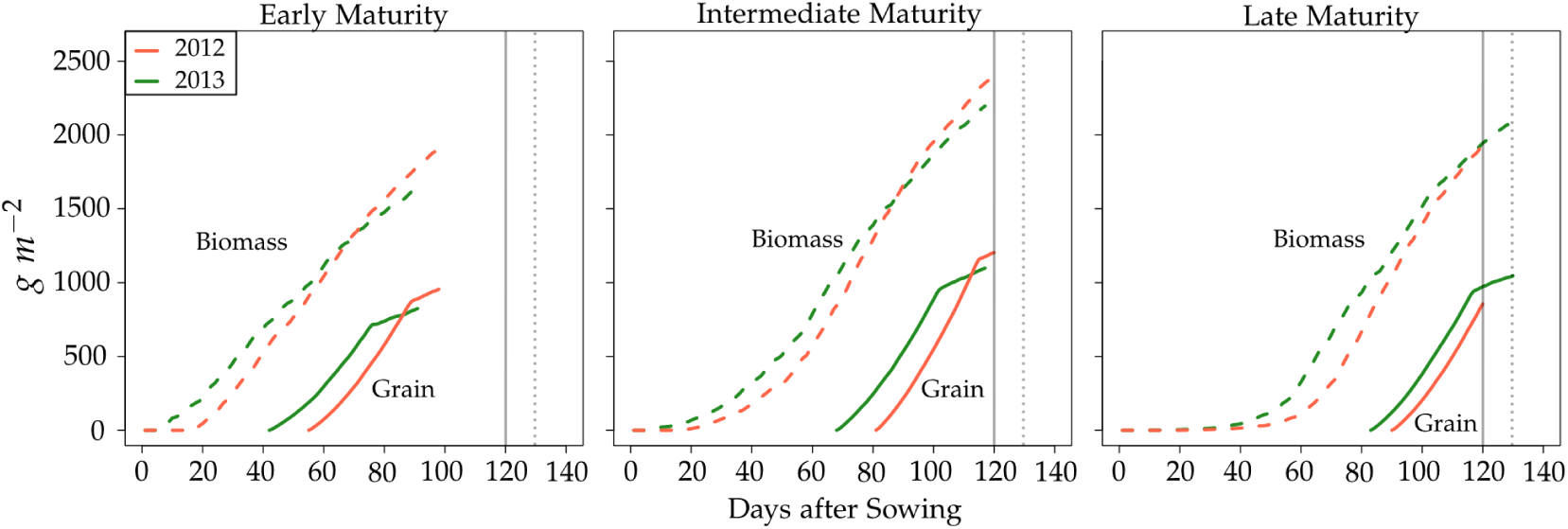
Simulated development of total biomass and grain yield. The early, intermediate and late maturing genotypes had a total leaf number (TLN) of 6, 14.5 and 23, respectively. The values for the other three traits were 750 for AM, 1.6 for SRE and 1150 for MTU and in common for all genotypes. The full and dotted vertical lines indicate the end of the 2012 and 2013 growing season, respectively.

The CGM can be viewed as a function *F* of the genotype specific inputs (the physiological traits) and the environment data

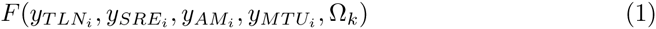

where 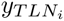 etc. are the values of the physiological traits observed for the *i^th^* genotype and the weather and management data of environment *k* are represented as Ω*_k_*. To simplify notation, we will henceforth use *F*(·)*_ik_* to represent the CGM and its inputs for genotype *i* in environment *k*.

## Approximate Bayesian Computation (ABC)

ABC replaces likelihood computation with a simulation step (Tavare et al. [42]). An integral component of any ABC algorithm is therefore the simulation model operator Model 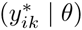 which generates simulated data 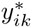 given parameters *θ*. In our proof of concept study, the crop growth model *F*(·)*_ik_* represents the deterministic component of Model 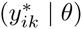, to which a Gaussian noise variable distributed as 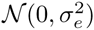 is added as a stochastic component. If Model 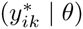 is fully deterministic, the distribution sampled with the ABC algorithm will not converge to the true posterior distribution when the tolerance for the distance between the simulated and observed data goes to zero (Sadegh and Vrugt [48]).

The weather and management data Ω*_k_* was assumed to be known, the physiological traits, however, were unknown and modeled as linear functions of the trait specific marker effects

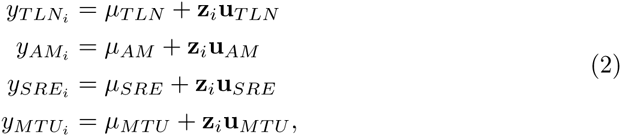

where **z***_i_* is the genotype vector of the observed biallelic single nucleotide polymorphism (SNP) markers of genotype *i*, *μ_TLN_* etc. denote the intercepts and **u***_TLN_* etc. the marker effects. For brevity, we will use *θ* to denote the joint parameter vector [*μ_TLN_*, …, *μ_MTU_*, **u***_TLN_*, …, **u***_MTU_*].

We used independent Normal distribution priors for all components of *θ*. The prior for *μ_TLN_* was 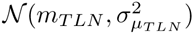. To simulate imperfect prior information, we drew the prior mean *m_TLN_* from a Uniform distribution over the interval 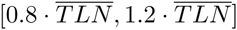, where 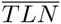 is the observed population mean of TLN. The average difference between *m_TLN_* and 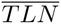 then is 10% of the latter value. The prior variance 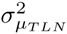, which represents the prior uncertainty, was equal to 2.25^2^. The prior means of AM, SRE and MTU were obtained accordingly and the prior variances 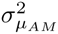, 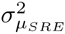 and 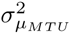 were 150^2^, 0.3^2^ and 225^2^, respectively.

The prior for the marker effects **u***_TLN_* was 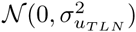, which corresponds to the *BayesC* prior (Habier et al. [58]). In BayesC, the prior variance of marker effects 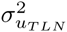, which introduces shrinkage, is the same across markers. For simplicity, we set this variance to a constant value and did not attempt to estimate it. Also in this case we simulated imperfect information by drawing the value of 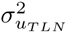 from a Uniform distribution over the interval [0.8 · *var*(*TLN*)/*M*, 1.2 · *var*(*TLN*)/*M*], where *M* is the number of markers and *var*(*TLN*) the observed population variance of TLN. The prior variances of marker effects of the other traits were obtained accordingly.

The value of 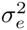, the variance of the Gaussian noise variable that is part of the model operator Model 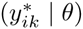, was drawn from a Uniform distribution over the interval [0.8 · *v_e_*, 1.2 · *v_e_*], where *v_e_* is the residual variance component of the phenotypic grain yield values used to fit the model.

Algorithm 1 in Table 1 shows pseudocode for the ABC rejection sampling algorithm we used. As distance measure between the simulated and observed data we used the Euclidean distance. The tolerance level *ϵ* for the distance between the simulated and observed data was tuned in a preliminary run of the algorithm to result in an acceptance rate of approximately 1 · 10^−6^. The number of posterior samples drawn was 100. We will refer to this ABC based WGP method that incorporates the CGM as CGM-WGP. The *CGM-WGP* algorithm was implemented as a C routine integrated with the R software environment (R Core Team [59]).

**Table 1.**
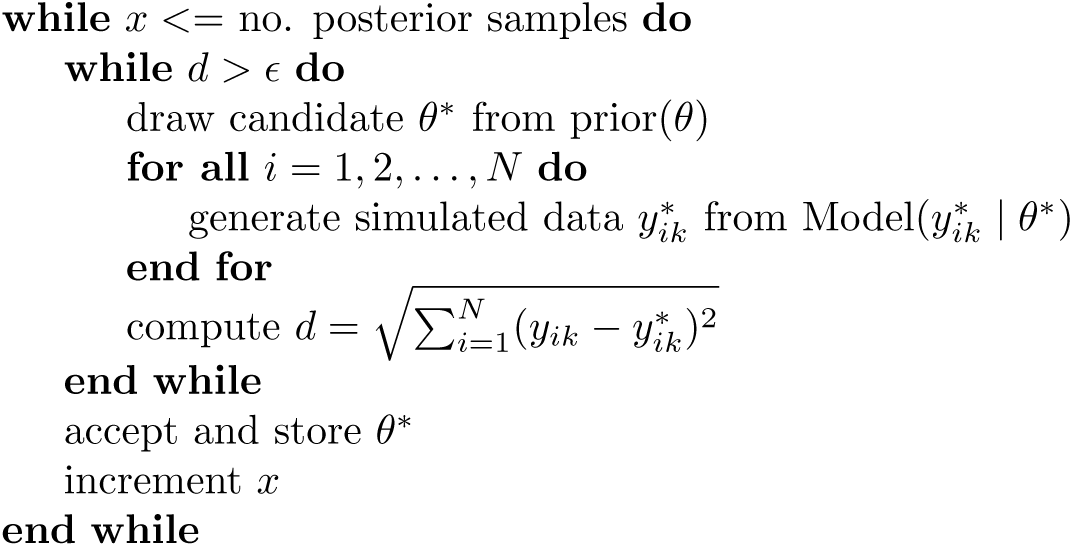
Pseudocode of ABC rejection sampling algorithm

### Synthetic data set

To test the performance of CGM-WGP, we created a biparental population of 1,550 doubled haploid (DH) inbred lines in silico. The genome consisted of a single chromosome of 1.5 Morgan length. The genotypes of the DH lines were generated by simulating meiosis events with the software package hypred (Technow [60]) according to the Haldane mapping function. On the chromosome, we equidistantly placed 140 informative SNP markers. A random subset of 40 of these markers were assigned to be QTL with additive effects on either TLN, AM, SRE or MTU. Each physiological trait was controlled by 10 of the 40 QTL, which were later removed from the set of observed markers available for analysis.

Basic ABC rejection sampling algorithm to sample from the approximate posterior distribution of *θ*.

The additive substitution effects of the QTL were drawn from a Standard Normal distribution. Raw genetic scores for each physiological trait were computed by summing the QTL effects according to the QTL genotypes of each DH line. These raw scores were subsequently re-scaled linearly to the aforementioned value ranges. Finally, phenotypic grain yield values were created as

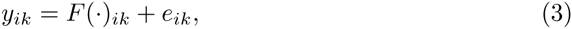

where *e_ik_* is a Gaussian noise variable with mean zero and variance *v_e_*. The value of *v_e_* was chosen such that the within-environment heritability of *y_ik_* was equal to 0.85. We generated 50 synthetic data sets by repeating the whole process.

### Estimation, prediction and testing procedure

The models were fitted using *N* = 50 randomly chosen DH lines as an estimation set. The remaining 1500 DH lines were used for testing model performance. Separate models were fitted using the 2012 and the 2013 grain yield data of the estimation set lines. The environment from which data for fitting the model was used will be referred to as *estimation environment*. Parameter estimates from each estimation environment were subsequently used to predict performance of the lines in the test set in both environments. Predictions for the same environment as the estimation environment will be referred to as *observed environment predictions* (e.g., predictions for 2012 with models fitted with 2012 data). Predictions for an environment from which no data were used in fitting the model will be referred to as *new environment predictions* (e.g., predictions for 2013 with models fitted with 2012 data).

As a point estimate for predicted grain yield performance in a specific environment, we used the mean of the posterior predictive distribution for the DH line in question. The posterior predictive distribution was obtained by evaluating *F*(·)*_ik_* over the accepted *θ* samples, using the weather and management data Ω*_k_* pertaining to that environment.

Prediction accuracy was computed as the Pearson correlation between predicted and true performance in the environment for which the prediction was made. The true grain yield performance was obtained by computing *F*(·)*_ik_* with the true values of the physiological traits.

As a performance benchmark we used genomic best linear unbiased prediction (GBLUP, Meuwissen et al. [1]). The model is

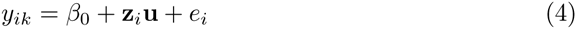

where *β*_0_ is the intercept, **u** the vector of marker effects and *e_i_* a residual. As before, **z***_i_* denotes the marker genotype vector. The GBLUP model was fitted with the R package rrBLUP (Endelman [61]). GBLUP and BayesC are comparable in their shrinkage behavior because both use a constant variance across markers. For GBLUP, predicted values were computed according to eq. (4) as *β*_0_ + **z***_i_***u**. Note that because the conventional GBLUP model does not utilize information about the environment for which predictions are made, observed and new environment predictions are identical.

## Results and Discussion

### Predicting performance in observed environments

The accuracy of observed environment predictions achieved by CGM-WGP was considerably larger than that of the benchmark method GBLUP in both environments (Table 2, Fig. 3, Fig. S2). This superiority of CGM-WGP over GBLUP can be explained by the presence of non-additive gene effects which cannot be captured fully by the latter. In the example scenario we studied, the non-additive gene effects on grain yield are a result of nonlinear functional relationships between the physiological traits and grain yield, which was particularly pronounced for TLN (Fig. 4).

**Table 2.** Accuracy of grain yield predictions of DH lines in the test set

| Estimation Env. | Prediction Env. | CGM-WGP | GBLUP |
| --- | --- | --- | --- |
| 2012 | 2012 | 0.77 | 0.54 |
|  | 2013 | 0.48 | 0.10 |
| 2013 | 2012 | 0.42 | 0.08 |
|  | 2013 | 0.75 | 0.62 |

Prediction accuracy for grain yield of DH lines in the test set, averaged over 50 replications. All differences within a row are statistically significant at a significance level of $< 0.005$ .

**Figure 3.**
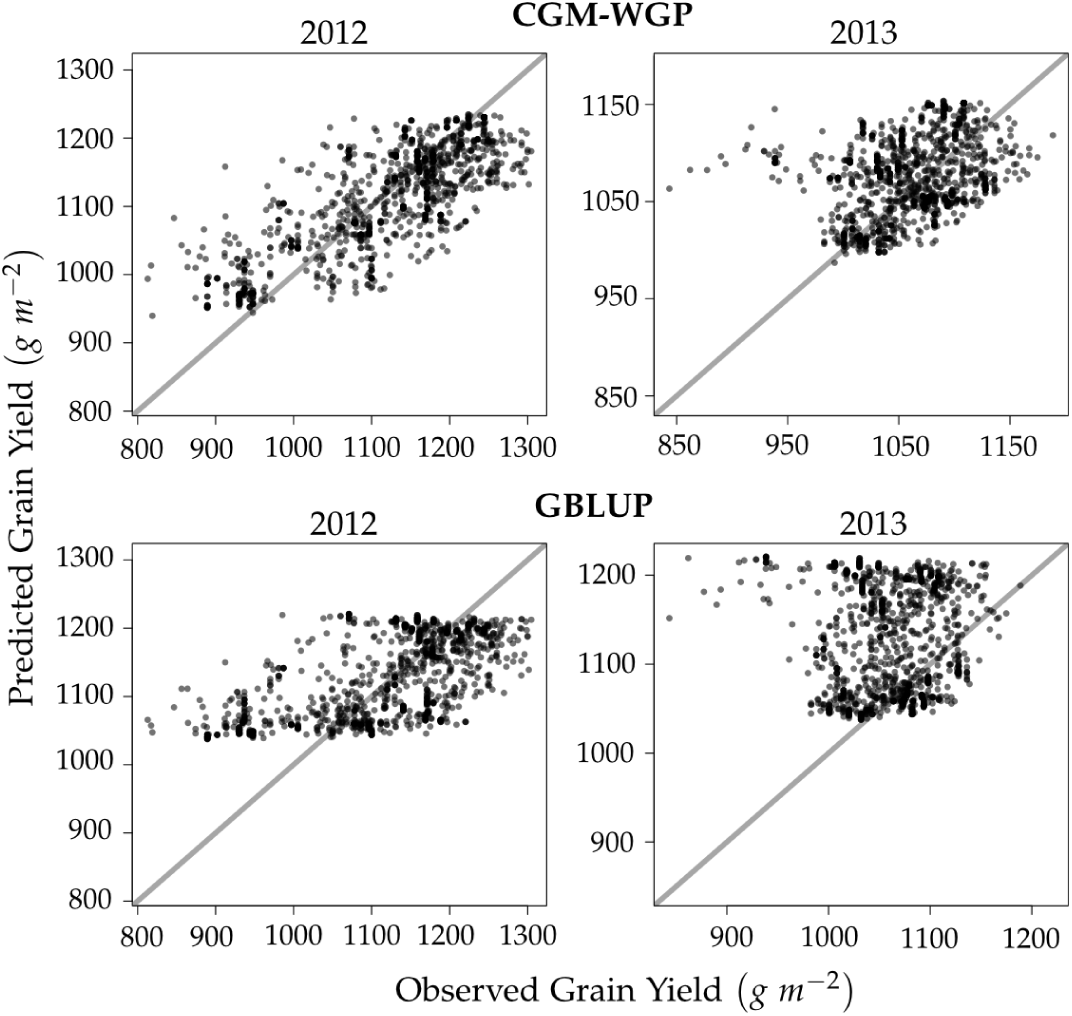
**Predicted vs. observed grain yield** of 1500 DH lines in testing set for prediction methods CGM-WGP (top row) and GBLUP (bottom row). The estimation environment was 2012. Results shown are from a representative example data set. In this example, the accuracy for observed environment predictions was 0.83 (CGM-WGP) and 0.69 (GBLUP). For new environment predictions it was 0.39 (CGM-WGP) and 0.11 (GBLUP).

**Figure 4.**
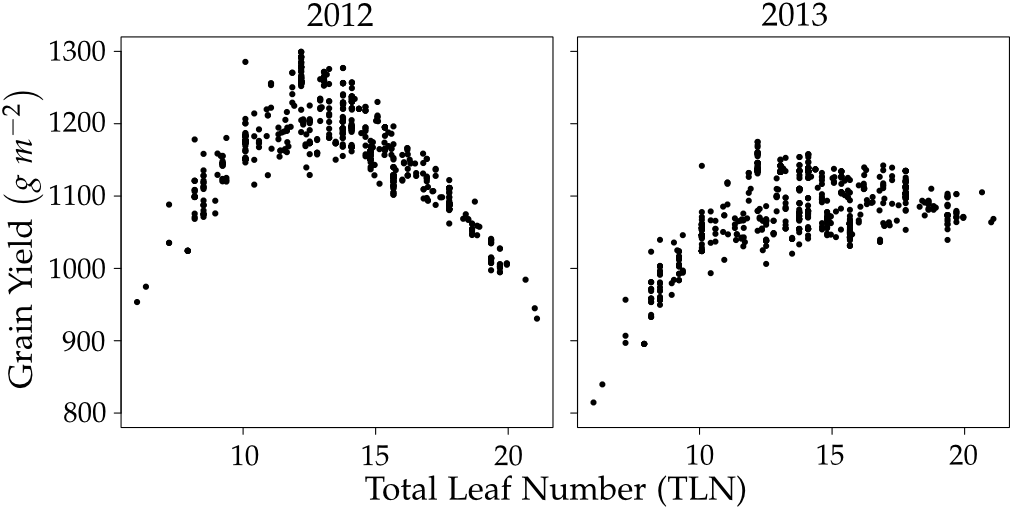
Relationship between total leaf number (TLN) and grain yield. Results shown are from a representative example data set.

TLN is closely related with the maturity rating of genotypes (Muchow et al. [50]). The higher it is, the later the onset of the reproductive phase and the later the maturity. Late genotypes have a higher yield potential than earlier genotypes because of a greater leaf area (Fig. S1). However, if the growing season is too short, they cannot realize this yield potential because of their slower development and later onset of the generative phase (Fig. 2). Very early genotypes on the other hand, have a low leaf area and do not make use of the full growing season. As a consequence, their realized yield is low, too. The relationship between TLN and grain yield therefore follows an optimum curve (Fig. 4). This was particularly pronounced in 2012, which had the shorter growing season and therfore penalized the late maturing genotypes more. The more decidedly nonlinear relationship between grain yield and TLN in 2012 also explains why the difference in prediction accuracy between CGM-WGP and GBLUP was greater in this season than in 2013 (0.23 points in 2012 compared to 0.13 points in 2013, on average).

Prediction accuracy for grain yield of DH lines in the test set, averaged over 50 replications. All differences within a row are statistically significant at a significance level of < 0.005.

The scenario we studied is an example of a particular case of epistasis, which might be called *biological epistasis*, that can arise even if the gene effects on the physiological component traits underlying the final trait of interest (grain yield in our case) are purely additive (Holland [33]). We accounted for nonlinear functional relationships among traits with the CGM. This enabled us to capture biological epistasis through simple linear models relating marker genotypes to the unobserved underlying physiological traits. Previously developed WGP models attempted to capture epistasis by directly fitting nonlinear marker effects to the final trait of interest (e.g., Xu [62]; Sun et al. [63]; Howard et al. [64]). While these models showed some promise, they have not been adopted by practitioners on a larger scale. By combining statistics with biological insights captured by CGMs, CGM-WGP takes a fundamentally different approach and presents a potentially powerful alternative to purely statistical WGP models.

### Predicting performance in new environments

New environment prediction accuracy was considerably lower than observed environment prediction accuracy, for both prediction methods (Table 2, Fig. 3, Fig. S2). The average prediction accuracy for performance in 2012 when using the 2013 estimation environment was 54% (CGM-WGP) and 15% (GBLUP) of the respective prediction accuracy achieved when using the 2012 estimation environment. The corresponding values for the accuracy of predicting performance in 2013 were 64% (CGM-WGP) and 16% (GBLUP). Thus, CGM-WGP still delivered a decent accuracy for predicting performance in new environments, while GBLUP largely failed in this task. The prediction accuracy of GBLUP was in fact negative, and sometimes strongly so, for close to 50% of the synthetic data sets (Fig. S2). For CGM-WGP negative accuracies were observed in only 14% (2012) and 4% (2013) of the cases.

The rank correlation between true performance in 2012 and 2013 was only 0.54 (averaged over 50 synthetic data sets), which indicated the presence of considerable G×E interactions, including changes in rank (Fig. S3). The interaction between the environment and TLN again explains the occurance of G×E to a large degree. In the shorter 2012 season, the late maturing genotypes cannot realize their growth and yield potential and are outperformed by the genotypes with early and intermediate maturity (Fig. 2 and Fig. 4). In the 10 day longer growing season of 2013, however, the late maturing genotypes can realize their greater yield potential better and outperform the early maturing genotypes and have a similar performance as genotypes with intermediate maturity. This dynamic leads to cross over G×E interactions between the 2012 and 2013 environments.

That new environment prediction under the presence of G×E interaction is considerably less accurate than observed environment prediction was expected and already observed in other studies (Resende et al. [65], Windhausen et al. [11]). It is encouraging that the reduction in accuracy for CGM-WGP was considerably less severe than for the conventional benchmark method GBLUP because this indicates that the former method did succeed in predicting G×E interactions to some degree.

Predicting G×E interactions in new environments for which no yield data are available, requires WGP models that link genetic effects (e.g., marker effects) with information that characterizes the environments. Jarquín et al. [40] accomplished this by fitting statistical interactions between markers and environmental covariates. A similar approach was taken by Heslot et al. [41], who in addition used a CGM to extract stress covariates from a large set of environmental variables. CGM-WGP takes this approach a step further by making the CGM and the environmental data that inform it, an integral part of the estimation procedure.

Nonetheless, while novel prediction methods might succeed in narrowing the gap between new and observed environment prediction, the former should always be expected to be less accurate than the latter. Field testing should therefore be performed in environments of particular importance for a breeding program to achieve the maximum attainable prediction accuracy for these. The same applies for target environments in which G×E interaction effects are expected to be particularly strong. CGMs can help to identify such environments and to inform experimental design and utilization of managed environments (Messina et al. [27], Messina et al. [29]). However, the range of the target population of environments of modern plant breeding programs is much too large for yield testing across the whole breadth (Cooper et al. [2]). Predicting performance in new environments will therefore always be required and novel methods like CGM-WGP are anticipated to be instrumental for enabling and enhancing success in this particularly daunting task.

### Further developments

#### More sophisticated CGMs

For this first proof of concept study, we assumed that the CGM used in the estimation process fully represented the systematic component of the data generating process, besides the random noise. This was clearly a “best case scenario”. However, decades of crop growth modeling research have provided the know-how necessary to approximate real crop development to any desired degree of accuracy (Keating et al. [17]), Renton [66], Hammer et al. [30]). Advanced CGMs such as *APSIM* (Keating et al. [17]), for example, model functional relationships between various crop parameters and external factors such as water and nutrient availability, soil properties as well as weed, insect and pathogen pressure. Thus, tools are principally available for applying CGM-WGP in more complex scenarios than the one addressed in this study.

#### Stochastic CGMs

There are examples of the use of fully deterministic model operators in ABC (Toni et al. [67], Liepe et al. [68]). However, with fully deterministic model operators the sampled distribution would not converge to the true posterior when the tolerance level *ϵ* goes to zero (Sadegh and Vrugt [48]) and instead reduce to a point mass over those parameter values that can reproduce the data. The CGM we used was fully deterministic. We therefore followed the example of Sadegh and Vrugt [48], who constructed a stochastic model operator by adding a random noise variable, with the same probabilistic properties as assumed for the residual component of the phenotype, to the deterministic functional model. A more elegant and possibly superior solution, however, would be to integrate stochastic processes directly into the CGM. While the vast majority of CGMs are deterministic (Keating et al. [17], van Ittersum et al. [16]), there are examples of stochastic CGMs (Brun et al. [69]). In addition to incorporating inherently stochastic processes of development (Curry et al. [70]), stochastic CGMs could also serve to account for uncertainty in the parameters of the functional equations comprising the model (Wallach et al. [71]).

#### Advanced ABC algorithms

For this proof of concept study we used the basic ABC rejection sampling algorithm (Tavare et al. [42], Pritchard et al. [43]). Considerable methodology related advances have been made, however, over the last decade that have led to algorithms with improved computational efficiency. Of particular interest here are population or sequential Monte Carlo algorithms, which are based on importance sampling (Sisson et al. [72], Toni et al. [67], Peters et al. [73]). These algorithms can dramatically increase acceptance rates without compromising on the tolerance levels. They achieve this by sampling from a sequence of intermediate proposal distributions of increasing similarity to the target distribution. Unfortunately, importance sampling fails when the number of parameters gets large, because then the importance weights tend to concentrate on very few samples, which leads to an extremely low effective sample size (Bengtsson et al. [74]). In the context of sequential Monte Carlo, this is known as particle depletion and was addressed by Peters et al. [73]. We implemented their approach, but were not able to overcome the problem of particle depletion. The number of parameters we estimated was 404 (100 marker effects per physiological trait plus an intercept), which seems well beyond the dimensionality range for importance sampling (Bengtsson et al. [74]).

Another interesting development is *MCMC-ABC*, which incorporates ABC with the Metropolis-Hastings algorithm (Marjoram et al. [75]). MCMC-ABC should result in high acceptance rates if the sampler moves into parameter regions of high posterior probability. However Metropolis-Hastings sampling too can be inefficient when the parameter space is of high dimension.

The greatest computational advantage of the original ABC rejection algorithm over Monte Carlo based ABC methods is that it generates independent samples and therefore readily lends itself to “embarrassingly” parallel computation (Marjoram et al. [75]). The computation time thus scales linearly to the number of processors available. Using the ABC rejection algorithm therefore allowed us to fully leverage the high performance computing cluster of DuPont Pioneer. In the era of cloud computing (Buyya et al. [76]), high performance computing environments are readily available to practitioners and scientists in both public and private sectors. Generality, scalability to parallel computations, and ease of implementation make the basic rejection sampler a viable alternative to more sophisticated approaches.

#### Using prior information

We used mildly informative prior distributions, the parameters of which were derived from the population means and variances of the physiological traits. In practice, the required prior information must be obtained from extraneous sources, such as past experiments or from the literature (Brun et al. [69]). Such information is imperfect and only partially matches the true population parameters of the population in question. We determined the prior parameters from the population itself, but perturbed them considerably to simulate erroneous prior information. Specifically, the average relative discrepancy (bias) between the prior parameter used and the true population parameter was 10%. When we increased the relative discrepancy to 25% (i.e., a maximum discrepancy of 50%), prediction accuracy dropped somewhat (Table S1). The reduction was only slight for observed environment prediction but more pronounced for new environment prediction. However, CGM-WGP was still considerably more accurate than the benchmark GBLUP. Thus, CGM-WGP seems to be relatively robust to moderate prior miss specification, as long as the value range supported by the prior distribution is not out of scope. In the ideal case of no prior bias, on the other hand, new and observed environment prediction accuracy increased slightly as compared to a bias of 10%.

In contrast to the complex trait of interest, component physiological traits may be realistically modeled based on a relatively simple genetic architecture, and for such traits, QTL explaining a sizable proportion of genetic variance can be mapped and characterized (Reymond et al. [77]; Bogard et al. [78]; Yin et al. [79]; Welcker et al. [80]; Tardieu et al. [81]). In fact, such component trait QTL have been successfully used to parametrize CGMs for studying genotype dependent response to environmental conditions (Chenu et al. [28]; Messina et al. [29] Bogard et al. [78]; Chenu et al. [82] Yin et al. [79]). Knowledge about the location and effect of such QTL, or of transgenes (Dong et al. [83]; Guo et al. [84]; Habben et al. [85]), could be incorporated as an additional source of prior information. Then, instead of estimating marker effects for the whole genome, CGM-WGP could focus on regions of particular importance, which reduces the dimensionality of the parameter space dramatically. Thus prior knowledge can be leveraged for improving prediction accuracy and computational efficiency of CGM-WGP.

#### Other applications

The idea of incorporating biological insights into WGP models is not limited to CGMs. Plant metabolites are chemical compounds produced as intermediate or end products of biochemical pathways. They are seen as potential bridges between genotypes and phenotypes of plants (Keurentjes [86]) and are therefore of particular interest in plant breeding (Fernie and Schauer [87]). Metabolic networks model the interrelationships between genes, intermediate metabolites and end products through biochemistry pathways (Schuster et al. [88]). Elaborate metabolic network models are available today that allow studying and simulating complex biochemical processes related to crop properties, such as flowering time, seed growth, nitrogen use efficiency and biomass composition (Dong et al. [83]; Pilalis et al. [89]; Simons et al. [90], Saha et al. [91]). Liepe et al. [47] demonstrated how ABC can be used for parameter estimation with metabolic and other biochemical networks. Using the principles outlined here for CGM-WGP, metabolic networks might add valuable biological information for the purpose of WGP, too.

Despite ever increasing sample sizes and marker densities, most of the genetic variance of complex traits remains unaccounted for in genome-wide association studies (Maher [92]). Marjoram et al. [49] argued that signal detection power could be increased by augmenting the purely statistical association models used thus far with biological knowledge. They demonstrated their approach by using ABC for incorporating gene regulatory networks into their analysis. Here we showed that the same principle can be applied to WGP by using ABC for integrating a CGM in the estimation of whole genome marker effects. Yield is a product of plant genetics and physiology, the environment and crop management and integrating information pertaining to these components will ultimately enable us to better predict it [93]. While this study is only a first step and many questions remain, we conclude that CGM-WGP presents a promising and novel path forward towards a new class of WGP models that leverage previously unused sources of knowledge and thereby increase prediction accuracy in settings that have proved challenging for plant breeding and applied genetics.

## Supporting Information

**Table S1.**Accuracy of grain yield predictions of test DH lines with increased bias in prior hyperparameters

**Figure S1.** Simulated development of total and senescent leaf area. The early, intermediate and late maturing genotypes had a total leaf number (TLN) of 6, 14.5 and 23, respectively. The values for the other three traits were 750 for AM, 1.6 for SRE and 1150 for MTU and in common for all genotypes. The full and dotted vertical lines indicate the end of the 2012 and 2013 growing season, respectively.

**Figure S2.** CGM-WGP vs. GBLUP prediction accuracy in 50 synthetic data sets.

**Figure S3.** Distribution of simulated grain yield in 2012 and 2013 environments. The grey lines indicate the performance of specific genotypes in both environments. Data shown is from a representative example replication.

**Figure S4.** Relationship between physiological traits and total grain yield. Data shown are a random sample of 1000 genotypes from a representative example replication.

## References

1. Meuwissen THE, Hayes BJ, Goddard ME (2001) Prediction of total genetic value using genome-wide dense marker maps. Genetics 157: 1819–1829.

2. Cooper M, Messina CD, Podlich D, Totir LR, Baumgarten A, et al. (2014) Predicting the future of plant breeding: complementing empirical evaluation with genetic prediction. Crop and Pasture Sci 64: 311–336.

3. Yang W, Tempelman RJ (2012) A Bayesian antedependence model for whole genome prediction. Genetics 190: 1491–1501.

4. Heslot N, Yang HP, Sorrells ME, Jannink JL (2012) Genomic selection in plant breeding: a comparison of models. Crop Sci 52: 146–160.

5. Kärkkäinen HP, Sillanpää MJ (2012) Back to basics for Bayesian model building in genomic selection. Genetics 191: 969–987.

6. Technow F, Melchinger AE (2013) Genomic prediction of dichotomous traits with Bayesian logistic models. Theor Appl Genet 126: 1133–1143.

7. Meuwissen T, Goddard M (2010) Accurate prediction of genetic values for complex traits by whole-genome resequencing. Genetics 185: 623–631.

8. Erbe M, Hayes BJ, Matukumalli LK, Goswami S, Bowman PJ, et al. (2012) Improving accuracy of genomic predictions within and between dairy cattle breeds with imputed high-density single nucleotide polymorphism panels. J Dairy Sci 95: 4114–4129.

9. Ober U, Ayroles JF, Stone EA, Richards S, Zhu D, et al. (2012) Using whole-genome sequence data to predict quantitative trait phenotypes in Drosophila melanogaster. PLoS Genet 8: e1002685.

10. Rincent R, Laloe D, Nicolas S, Altmann T, Brunel D, et al. (2012) Maximizing the reliability of genomic selection by optimizing the calibration set of reference individuals: comparison of methods in two diverse groups of maize. Genetics 192: 715–728.

11. Windhausen VS, Atlin GN, Hickey JM, Crossa J, Jannink JL, et al. (2012) Effectiveness of genomic prediction of maize hybrid performance in different breeding populations and environments. G3 2: 1427–1436.

12. Technow F, Bürger A, Melchinger AE (2013) Genomic prediction of northern corn leaf blight resistance in maize with combined or separated training sets for heterotic groups. G3 3: 197–203.

13. Hickey JM, Dreisigacker S, Crossa J, Hearne S, Babu R, et al. (2014) Evaluation of genomic selection training population designs and genotyping strategies in plant breeding programs using simulation. Crop Sci in press: doi:10.2135/cropsci2013.03.0195.

14. Daetwyler HD, Pong-Wong R, Villanueva B, Woolliams JA (2010) The impact of genetic architecture on genome-wide evaluation methods. Genetics 185: 1021–1031.

15. Habier D, Fernando RL, Garrick DJ (2013) Genomic-BLUP decoded: a look into the black box of genomic prediction. Genetics 194: 597–607.

16. van Ittersum MK, Leffelaar PA, Van Keulen H, Kropff MJ, Bastiaans L, et al. (2003) On approaches and applications of the Wageningen crop models. Eur J Agron 18: 201–234.

17. Keating BA, Carberry PS, Hammeer GL, Probert ME, Robertson MJ, et al. (2003) An overview of APSIM, a model designed for farming systems simulation. Eur J Agron 18: 267–288.

18. Cooper M, van Eeuwijk FA, Hammer GL, Podlich D, Messina C (2009) Modeling QTL for complex traits: detection and context for plant breeding. Curr Opin Plant Biol 12: 231–240.

19. Hammer G, Cooper M, Tardieu F, Welch S, Walsh B, et al. (2006) Models for navigating biological complexity in breeding improved crop plants. Trends Plant Sci 11: 587–593.

20. Passioura JB (1983) Roots and drought resistance. Agr Water Manage 7: 265–280.

21. Yin X, Struik PC, Kropff MJ (2004) Role of crop physiology in predicting gene-to-phenotype relationships. Trends Plant Sci 9: 426–432.

22. Chapman S, Cooper M, Hammer G, Butler D (2000) Genotype by environment interactions affecting grain sorghum. ii. frequencies of different seasonal patterns of drought stress are related to location effects on hybrid yields. Aust J Agric Res 51: 209–222.

23. Löffler CM, Wei J, Fast T, Gogerty J, Langton S, et al. (2005) Classification of maize environments using crop simulation and geographic information systems. Crop Sci 45: 1708–1716.

24. Hammer G, Dong Z, McLean G, Doherty A, Messina C, et al. (2009) Can changes in canopy and/or root system architecture explain historical maize yield trends in the U.S. corn belt? Crop Sci 49: 299–312.

25. Chapman S, Cooper M, Podlich D, Hammer G (2003) Evaluating plant breeding strategies by simulating gene action and dryland environment effects. Agron J 95: 99–113.

26. Messina C, Hammer G, Dong Z, Podlich D, Cooper M (2009) Chapter 10 - Modelling crop improvement in a G×E×M framework via gene-trait-phenotype relationships. In: Sadras V, Calderini D, editors, Crop Physiology, San Diego: Academic Press. pp. 235–581.

27. Messina CD, Podlich D, Dong Z, Samples M, Cooper M (2011) Yield-trait performance landscapes: from theory to application in breeding maize for drought tolerance. J Exp Bot 62: 855–868.

28. Chenu K, Chapman SC, Hammer GL, McLean G, Salah HB, et al. (2008) Short-term responses of leaf growth rate to water deficit scale up to whole-plant and crop levels: an integrated modelling approach in maize. Plant Cell Environ 31: 378–391.

29. Messina CD, Jones JW, Boote KJ, Vallejos CE (2006) A gene-based model to simulate soybean development and yield responses to environment. Crop Sci 46: 456–466.

30. Hammer GL, van Oosterom E, McLean G, Chapman SC, Broad I, et al. (2010) Adapting APSIM to model the physiology and genetics of complex adaptive traits in field crops. J Exp Bot 61: 2185–2202.

31. Wolf DP, Hallauer AR (1997) Triple testcross analysis to detect epistasis in maize. Crop Sci 37: 736–770.

32. Eta-Ndu JT, Openshaw SJ (1999) Epistasis for grain yield in two F_2_ populations of maize. Crop Sci 39: 346–352.

33. Holland JB (2001) Epistasis and plant breeding. In: Janick J, editor, Plant Breeding Reviews, Volume 21, Hoboken, NJ: John Wiley & Sons, Inc. pp. 27–92.

34. Cooper M, Chapman SC, Podlich D, Hammer G (2002) The GP problem: quantifying gene-to-phenotype relationships. In Silico Biol 2: 151–164.

35. Riedelsheimer C, Lisec J, Czedik-Eysenberg A, Sulpice R, Flis A, et al. (2012) Genome-wide association mapping of leaf metabolic profiles for dissecting complex traits in maize. Proc Natl Acad Sci 109: 8872–8877.

36. Richey FD (1942) Mock-dominance and hybrid vigor. Science 96: 280–281.

37. Melchinger AE, Singh M, Link W, Utz H, von Kittlitz E (1994) Heterosis and gene effects of multiplicative characters: theoretical relationships and experimental results from Vicia faba L. Theor Appl Genet 88: 343–348.

38. Schulz-Streeck T, Ogutu JO, Gordillo A, Karaman Z, Knaak C, et al. (2013) Genomic selection allowing for marker-by-environment interaction. Plant Breeding 132: 532–538.

39. Burgeño J, de los Campos G, Weigel K, Crossa J (2012) Genomic prediction of breeding values when modeling genotype x environment interaction using pedigree and dense molecular markers. Crop Sci 52: 702–719.

40. Jarquín D, Crossa J, Lacaze X, Du Cheyron P, Daucourt J, et al. (2014) A reaction norm model for genomic selection using high-dimensional genomic and environmental data. Theor Appl Genet 127: 595–607.

41. Heslot N, Akdemir D, Sorrells ME, Jannink JL (2014) Integrating environmental covariates and crop modeling into the genomic selection framework to predict genotype by environment interactions. Theor Appl Genet 127: 463–480.

42. Tavare S, Balding DJ, Griffiths RC, Donnelly P (1997) Inferring coalescence times from DNA sequence data. Genetics 145: 505–518.

43. Pritchard J, Seielstad M, Perez-Lezaun A, Feldman M (1999) Population growth of human Y chromosomes: A study of Y chromosome microsatellites. Mol Biol Evol 16: 1791–1798.

44. Csilléry K, Blum MG, Gaggiotti OE, François O (2010) Approximate Bayesian Computation (ABC) in practice. Trends Ecol Evol 25: 410–418.

45. Lopes JS, Beaumont MA (2010) ABC: A useful Bayesian tool for the analysis of population data. Infect Genet and Evol 10: 825–832.

46. Lawson Handley LJ, Estoup A, Evans DM, Thomas CE, Lombaert E, et al. (2011) Ecological genetics of invasive alien species. BioControl 56: 409–428.

47. Liepe J, Kirk P, Filippi S, Toni T, Barnes CP, et al. (2014) A framework for parameter estimation and model selection from experimental data in systems biology using approximate Bayesian computation. Nat Protoc 9: 439–456.

48. Sadegh M, Vrugt JA (2014) Approximate bayesian computation using Markov chain Monte Carlo simulation: DREAM(ABC). Water Resour Res 50: 6767–6787.

49. Marjoram P, Zubair A, Nuzhdin SV (2014) Post-GWAS: where next? More samples, more SNPs or more biology? Heredity 112: 79–88.

50. Muchow RC, R ST, Bennett JM (1990) Temperature and solar radiation effects on potential maize yield across locations. Agron J 82: 338–343.

51. Meghji MR, Dudley JW, Lambert RJ, Sprague GF (1984) Inbreeding depression, inbred and hybrid grain yields, and other traits of maize genotypes representing three eras. Crop Sci 24: 545–549.

52. Elings A (2000) Estimation of leaf area in tropical maize. Agron J 92: 436–444.

53. Muchow RC, Davis R (1988) Effect of nitrogen supply on the comparative productivity of maize and sorghum in a semi-arid tropical environment ii. radiation interception and biomass accumulation. Field Crop Res 18: 17–30.

54. McGarrahan JP, Dale RF (1984) A trend toward a longer grain-filling period for corn: a case study in Indiana. Agron J 76: 518–522.

55. Muchow R (1990) Effect of high temperature on grain-growth in field-grown maize. Field Crop Res 23: 145–158.

56. Nielsen RL, Thomison PR, Brown GA, Halter AL, Wells J, et al. (2002) Delayed planting effects on flowering and grain maturation of dent corn. Agron J 94: 549–558.

57. Neild, RE, Newman, JE. (1987) Growing season characteristics and requirements in the Corn Belt. Rep. NCH 40. Purdue Univ., West Lafayette, IN.

58. Habier D, Fernando R, Kizilkaya K, Garrick D (2011) Extension of the Bayesian alphabet for genomic selection. BMC Bioinformatics 12: 186.

59. R Core Team (2014) R: A Language and Environment for Statistical Computing. R Foundation for Statistical Computing, Vienna, Austria. URL <u>http://www.R-project.org/.</u>

60. Technow F (2013) hypred: Simulation of genomic data in applied genetics. R package version 0.4.

61. Endelman JB (2011) Ridge regression and other kernels for genomic selection with R package rrBLUP. Plant Genome 4: 250–255.

62. Xu S (2007) An empirical Bayes method for estimating epistatic effects of quantitative trait loci. Biometrics 63: 513–521.

63. Sun X, Ma P, Mumm RH (2012) Nonparametric method for genomics-based prediction of performance of quantitative traits involving epistasis in plant breeding. PLoS ONE 7: e50604.

64. Howard R, Carriquiry AL, Beavis WD (2014) Parametric and nonparametric statistical methods for genomic selection of traits with additive and epistatic genetic architectures. G3 4: 1027–1046.

65. Resende MF, Munoz P, Acosta JJ, Peter GF, Davis JM, et al. (2012) Accelerating the domestication of trees using genomic selection: accuracy of prediction models across ages and environments. New Phytol 193: 617–624.

66. Renton M (2011) How much detail and accuracy is required in plant growth sub-models to address questions about optimal management strategies in agricultural systems? AoB Plants 2011: plr006.

67. Toni T, Welch D, Strelkowa N, Ipsen A, Stumpf MP (2009) Approximate Bayesian computation scheme for parameter inference and model selection in dynamical systems. J R Soc Interface 6: 187–202.

68. Liepe J, Barnes C, Cule E, Erguler K, Kirk P, et al. (2010) ABC-SysBio approximate Bayesian computation in Python with GPU support. Bioinformatics 26: 1797–1799.

69. Brun F, Wallach D, Makowski D, Jones JW (2006) Working with dynamic crop models: Evaluation, analysis, parameterization, and applications. Amsterdam: Elsevier.

70. Curry GL, Feldman RM, Sharpe PJH (1978) Foundations of stochastic development. J Theor Biol 74: 397–410.

71. Wallach D, Keussayan N, Brun F, Lacroix B, Bergez JE (2012) Assessing the uncertainty when using a model to compare irrigation strategies. Agron J 104: 1274–1283.

72. Sisson SA, Fan Y, Tanaka MM (2007) Sequential Monte Carlo without likelihoods. Proc Nat Acad Sci 104: 1760–1765.

73. Peters GW, Fan Y, Sisson SA (2012) On sequential Monte Carlo, partial rejection control and approximate Bayesian computation. Stat Comput 22: 1209–1222.

74. Bengtsson T, Bickel P, Li B (2008) Curse-of-dimensionality revisited: Collapse of the particle filter in very large scale systems. In: IMS Collections Probability and Statistics: Essays in Honor of David A. Freedman, Institute of Mathematical Statistics, volume 2. pp. 316–334.

75. Marjoram P, Molitor J, Plagnol V, Tavaré S (2003) Markov chain Monte Carlo without likelihoods. Proc Natl Acad Sci 100: 15324–15328.

76. Buyya R, Yeo CS, Venugopal S (2008) Market-oriented cloud computing: vision, hype, and reality for delivering IT services as computing utilities. In: High Performance Computing and Communications, 2008. HPCC ’08. 10th IEEE International Conference on. pp. 5–13.

77. Reymond M, Muller B, Leonardi A, Charcosset A, Tardieu F (2003) Combining quantitative trait Loci analysis and an ecophysiological model to analyze the genetic variability of the responses of maize leaf growth to temperature and water deficit. Plant Physiol 131: 664–675.

78. Bogard M, Ravel C, Paux E, Bordes J, Balfourier F, et al. (2014) Predictions of heading date in bread wheat (Triticum aestivum L.) using QTL-based parameters of an ecophysiological model. J Exp Bot 10.1093/jxb/eru328.

79. Yin X, Struik PC, van Eeuwijk FA, Stam P, Tang J (2005) QTL analysis and QTL-based prediction of flowering phenology in recombinant inbred lines of barley. J Exp Bot 56: 967–976.

80. Welcker C, Boussuge B, Bencivenni C, Ribaut JM, Tardieu F (2007) Are source and sink strengths genetically linked in maize plants subjected to water deficit? A QTL study of the responses of leaf growth and of Anthesis-Silking Interval to water deficit. J Exp Bot 58: 339–349.

81. Tardieu F, Reymond M, Muller B, Granier C, Simonneau T, et al. (2005) Linking physiological and genetic analyses of the control of leaf growth under changing environmental conditions. Crop and Pasture Sci 56: 937–946.

82. Chenu K, Chapman SC, Tardieu F, McLean G, Welcker C, et al. (2009) Simulating the yield impacts of organ-level quantitative trait loci associated with drought response in maize: a “gene-to-phenotype” modeling approach. Genetics 183: 1507–1523.

83. Dong Z, Danilevskaya O, Abadie T, Messina C, Coles N, et al. (2012) A gene regulatory network model for floral transition of the shoot apex in maize and its dynamic modeling. PLoS ONE 7: e43450.

84. Guo M, Rupe MA, Wei J, Winkler C, Goncalves-Butruille M, et al. (2014) Maize ARGOS1 (ZAR1) transgenic alleles increase hybrid maize yield. J Exp Bot 65: 249–260.

85. Habben JE, Bao X, Bate NJ, DeBruin JL, Dolan D, et al. (2014) Transgenic alteration of ethylene biosynthesis increases grain yield in maize under field drought-stress conditions. Plant Biotechnol J 12: 685–693.

86. Keurentjes JJB (2009) Genetical metabolomics: closing in on phenotypes. Curr Opin Plant Biol 12: 223–230.

87. Fernie AR, Schauer N (2009) Metabolomics-assisted breeding: a viable option for crop improvement? Trends Genet 25: 39–48.

88. Schuster S, Fell DA, Dandekar T (2000) A general definition of metabolic pathways useful for systematic organization and analysis of complex metabolic networks. Nat Biotechnol 18: 326–332.

89. Pilalis E, Chatziioannou A, Thomasset B, Kolisis F (2011) An in silico compartmentalized metabolic model of brassica napus enables the systemic study of regulatory aspects of plant central metabolism. Biotechnol and Bioeng 108: 1673–1682.

90. Simons M, Saha R, Guillard L, Clément G, Armengaud P, et al. (2014) Nitrogen-use efficiency in maize (Zea mays L.): from ‘omics’ studies to metabolic modelling. J Exp Bot 65: 5657–5671.

91. Saha R, Suthers PF, Maranas CD (2011) Zea mays RS1563: A comprehensive genome-scale metabolic reconstruction of maize metabolism. PLoS ONE 6: e21784.

92. Maher B (2008) Personal genomes: The case of the missing heritability. Nature 456: 18–21.

93. Nature Genetics Editorial (2015) Growing access to phenotype data. Nat Genet 47: 99.

